# Dynamic phosphorylation of the *Plasmodium falciparum* Myosin A motor regulates motor speed and is important for asexual replication

**DOI:** 10.64898/2026.09.16.752031

**Authors:** Stephanie D. Nofal, Konstantinos Koussis, Juha Vahokoski, Inari Kursula, Justin E. Molloy, Michael J. Blackman, Christian Flueck, David A. Baker

## Abstract

Malaria parasite invasion of red blood cells is powered by Myosin A (MyoA), the actomyosin motor which is a core component of the glideosome. Our previous phosphoproteomic study identified a cyclic GMP-dependent phosphorylation site on MyoA at Ser19, suggesting that cyclic nucleotide signalling may directly regulate motor activity during invasion. However, the functional significance of this phosphorylation site remains unclear.

Here, we show that *Plasmodium falciparum* MyoA phosphorylation at Ser19 is dynamically regulated during the intraerythrocytic cycle, peaking in extracellular merozoites and rapidly declining following invasion. Using conditional depletion of MyoA, we show that MyoA is essential for red blood cell invasion but dispensable for gametocyte development. Conditional complementation with mutant versions of MyoA that either mimic (S19D) or ablate phosphorylation (S19A) reveal markedly impaired asexual parasite proliferation, highlighting the importance of precise regulation of MyoA phosphorylation at Ser19. *In vitro* motility assays using parasite-derived MyoA showed that ablating phosphorylation either pharmacologically or by introducing an S19A mutation, reduces motor speed. Conversely, introducing a phosphomimetic S19D mutation restores motility levels comparable to phosphorylated wild type MyoA. Collectively, these results establish MyoA Ser19 phosphorylation as a regulatory switch for actomyosin motor activity.

## Introduction

Malaria, caused by apicomplexan parasites of the genus *Plasmodium*, claims approximately 600,000 lives every year (WHO, 2025). The clinical manifestations of the disease arise due to repeated rounds of asexual replication of the parasite within red blood cells (RBCs) of the human host. After successful replication, the parasite undergoes a tightly regulated process called egress, whereby invasive merozoites are released from the infected RBC and go on to invade a new RBC to initiate the next cycle of blood stage development. Erythrocyte invasion is powered by the glideosome, a macromolecular complex localised between the parasite plasma membrane (PPM) and inner membrane complex (IMC). The glideosome consists of several components, including a class XIV actomyosin motor known as Myosin A (MyoA) (Pinder et al., 1998), an essential light chain (ELC) (Bookwalter et al., 2017; Green et al., 2017), a Myosin A-tail interacting protein (MTIP) (Bergman, 2003) known as ELC1 in the related apicomplexan parasite *Toxoplasma gondii* and the glideosome-associated proteins GAP40, GAP45 and GAP50 (Baum et al., 2005; Frénal et al., 2010; Gaskins et al., 2004). MTIP and ELC bind to the tail region of MyoA (Green et al., 2006) and recruit it to GAP45, which tethers the PPM to the IMC (Rees-Channer et al., 2006). This complex is then able to interact with the integral IMC proteins, GAP50 and GAP40, to form the assembled glideosome (Gaskins et al., 2004). Force generated by the motor translocates actin filaments rearwards towards the basal end of the parasite. These filaments are in turn tethered to adhesins on the surface of the parasite, which bind to receptors on the erythrocyte; allowing the parasite to glide across the surface of erythrocytes, wrap the host cell around itself in a process called deformation and invade (Yahata et al., 2021). The role of the glideosome in powering *Plasmodium falciparum* invasion has been demonstrated by several studies, whereby conditional deletion of either GAP45 (Perrin et al., 2018), MyoA (Robert-Paganin et al., 2019), ELC1 (Moussaoui et al., 2020) or more recently GAP40 (He et al., 2023) resulted in merozoites that were unable to deform or invade new erythrocytes.

During the final stages of merozoite development within the erythrocyte, cyclic GMP (cGMP), cyclic AMP (cAMP) and calcium-activated signalling pathways work in concert, resulting in the release of invasive merozoites. This includes the activation of the cGMP-dependent protein kinase (PKG), leading to the release of calcium and activation of calcium dependent-kinases required for egress, and finally the activation of the cAMP-dependent protein kinase (PKA) (Baker et al., 2017). Our earlier phosphoproteomic study identified cGMP-dependent phosphorylation of several components of the glideosome, including MyoA, GAP40 & GAP45; suggesting that signalling events downstream of PKG activation may be involved in regulating invasion (Alam et al., 2015). The most compelling PKG-dependent phosphosite is found on MyoA at serine 19 (Ser19), since phosphorylation of the equivalent residue in *Toxoplasma gondii* (Ser21) regulates the activity of the TgMyoA motor (Gaji et al., 2015; Tang et al., 2014). Moreover, a recent crystal structure of PfMyoA has revealed that phosphorylated Ser19 forms an electrostatic interaction with Lys764; a residue in the converter region of the motor (Robert-Paganin et al., 2019). This electrostatic interaction likely directly controls the speed and force of the motor, since motility assays using recombinant MyoA harbouring S19A or K764E mutations that block this interaction revealed a reduction in motor speed and ATPase activity (Robert-Paganin et al., 2019). Constitutive ectopic expression of the K764E mutant in *P. falciparum* MyoA knockout parasites led to a moderate growth defect due to slower internalisation of merozoites and an increased rate of merozoite ejection after internalisation of the parasite (Blake et al., 2020). In the related parasite, *P. berghei,* it was shown that the S19A mutation severely impacted mosquito to mouse transmission, likely due to the inability of sporozoites harbouring the mutation to colonise the salivary glands of mosquitoes (Ripp et al., 2022).

Despite several lines of evidence suggesting that phosphorylation of Ser19 modulates the molecular dynamics of the MyoA motor, this has yet to be shown in *P. falciparum* parasites. Furthermore, it is unclear whether timely dephosphorylation of this site is also important for motor function and parasite survival.

Here, we show in *P. falciparum* parasites that phosphorylation of MyoA at Ser19 occurs rapidly prior to egress, with maximal phosphorylation levels observed in extracellular merozoites, followed by dephosphorylation of this site shortly after invasion. We also demonstrate that this phosphorylation event is important, but not essential for asexual parasite growth since parasites harbouring mutations that block Ser19 phosphorylation are able to proliferate, albeit at a considerably reduced rate compared to wild type parasites. Importantly, simulating constitutive phosphorylation of MyoA by conditionally complementing the knockouts with MyoA harbouring an S19D phosphomimetic mutation is also detrimental to parasite proliferation, leading to a marked decrease in parasite growth. By performing *in vitro* motility assays using parasite-derived MyoA, we confirm that the S19A mutation reduces *in vitro* motor motility, while the S19D mutation restores motor motility to levels comparable to wild type phosphorylated MyoA. Considering that both S19A and S19D mutations are detrimental to parasite growth, despite the S19D mutation restoring the motor speed to that of phosphorylated MyoA, we show that timely phosphorylation of MyoA followed by its rapid dephosphorylation following invasion is important for asexual parasite growth. Finally, we demonstrate that by fine-tuning the degree of MyoA phosphorylation pharmacologically, we can regulate the speed of the motor, confirming the role of this phosphosite in modulating the activity of the actomyosin motor.

## Results

### Phosphorylation of MyoA Ser19 occurs rapidly and is likely important for asexual parasite proliferation

Two previous studies have shown that phosphorylation of MyoA at Ser19 in mature *P. falciparum* schizonts occurs in a PKG-dependent manner, since phosphorylation of this residue is drastically reduced in schizonts treated with the reversible PKG-specific inhibitor compound 2 (C2) (Alam et al., 2015; Flueck et al., 2019). To further investigate the dynamics of this phosphorylation event upon PKG activation in mature schizonts, we monitored the levels of MyoA phosphorylation in a MyoA-mCherry tagged line (Supplementary Fig.1) over time by western blot analysis using a MyoA Ser19 phospho-specific antibody (Alam et al., 2015) following removal of C2. To block parasite egress following C2 removal, the parasites were pre-treated with the irreversible cysteine protease inhibitor E64, which acts downstream of PKG activation (Hale et al., 2017; Salmon et al., 2001; Thomas et al., 2018) (Figure 1A). This revealed that MyoA Ser19 is rapidly phosphorylated following PKG activation, with maximal phosphorylation detected 14-18 minutes following release of the C2 block (Figure 1B), which corresponds to the average time taken for merozoites to egress following removal of C2 (Ressurreição et al., 2020).

**Figure 1.**
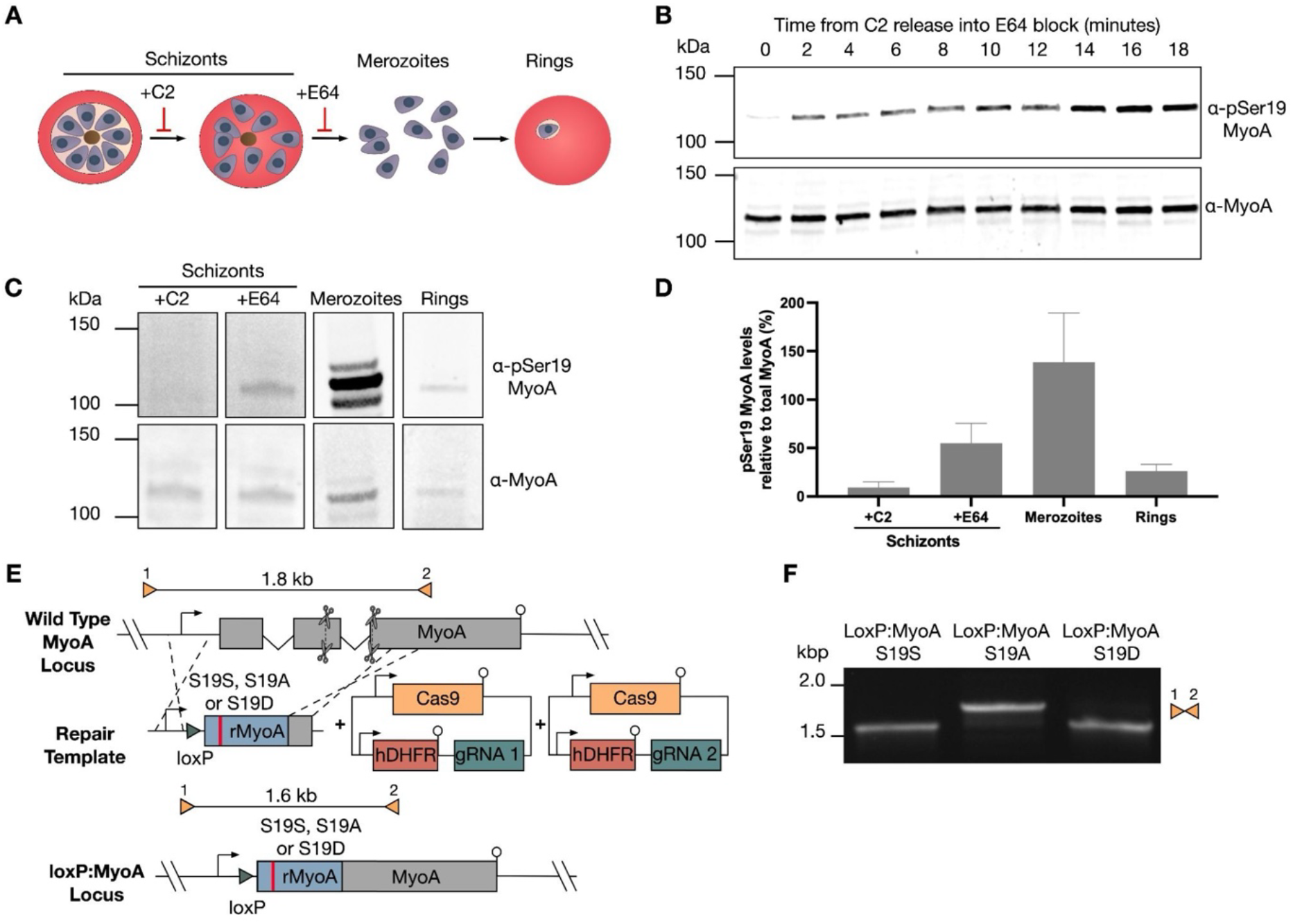
Dynamic phosphorylation of MyoA during the asexual intraerythrocytic cycle and generation of MyoA Ser19 mutant parasites. **A** Schematic showing where C2 and E64 act during the intraerythrocytic cycle to block parasite egress. C2 blocks egress prior to parasitophorous vacuole membrane rupture, whereas E64 blocks egress prior to erythrocyte membrane rupture. **B** Western blot analysis of MyoA Ser19 phosphorylation following the release of C2-arrested schizonts into E64-containing medium. **C** Western blot analysis of MyoA Ser19 phosphorylation and total MyoA across the parasite stages shown in Fig. 1A. **D** Densitometric quantification of MyoA Ser19 phosphorylation relative to total MyoA. Data presented are means from two independent experiments, with a representative western blot shown in Fig. 1C. Error bars represent standard deviation. **E** Schematic of the marker-free CRISPR/Cas9-mediated approach used to introduce the S19A, S19D or S19S (WT control) sequence into the endogenous *myoA* locus. Scissors indicate CRISPR/Cas9 cleavage sites. Coloured triangles represent primer pairs used for diagnostic PCR screens, together with the expected amplicon sizes of the non-integrated (1.8 kb) and integrated (1.6 kb) loci. **F** Diagnostic PCR analysis of parasite lines transfected with the S19S (WT control, S19A and S19D repair templates using the primer pairs shown in Fig. 1D.

To investigate whether any changes in phosphorylation of MyoA Ser19 could be observed during the events leading up to and including egress and invasion, the phospho-status of C2/E64-blocked schizonts, purified merozoites and newly formed rings at 0-4 hours post invasion were quantified by western blot. This revealed distinct changes in the phospho-status of MyoA as the parasites transition from schizonts to merozoites and then rings (Figure 1C), with a 2.5-fold increase in MyoA Ser19 phosphorylation in merozoites compared to E64-blocked schizonts, followed by a 5-fold reduction in rings (Figure 1D). These dramatic changes in phosphorylation over the course of egress and invasion point towards a regulatory role for this modification, with phosphorylation possibly activating the motor prior to invasion and dephosphorylation perhaps contributing to its inactivation or disassembly in early rings.

To determine if this phosphorylation site is essential for parasite proliferation, we attempted to modify the parasite’s MyoA locus by introducing mutations that would either block phosphorylation (MyoA S19A), mimic constitutive phosphorylation (MyoA S19D) of MyoA, or introduce a synonymous mutation, leaving the amino acid sequence unchanged (MyoA S19S) as a control (Figure 1E). PCR analysis revealed successful integration of the S19D and control repair templates (Figure 1F). However, after three attempts we were unable to integrate the S19A mutation, suggesting that phosphorylation of MyoA Ser19 is vital for asexual parasite proliferation. Although we successfully generated a parasite line harbouring the S19D mutation, we did not investigate its impact on parasite growth. Instead, we reasoned that a conditional mutagenesis strategy would offer a more rigorous approach to dissect the functional consequences of the S19D mutation, while avoiding potential adaptations that could arise over multiple replication cycles. This approach would also allow us to conditionally introduce the S19A mutation, which we were unable to generate through direct mutagenesis.

### MyoA is essential for asexual parasite invasion of erythrocytes

Since introducing the S19A mutation into the endogenous *myoa* gene proved unsuccessful, we aimed to develop a system that would allow for conditional expression of MyoA harbouring Ser19 mutations, while simultaneously deleting the endogenous copy. To achieve this, we first generated a parasite line in which we could conditionally knock out (cKO) the endogenous *myoa* gene using the dimerisable Cre recombinase (DiCre) system (Collins et al., 2013). The conditional knockout was engineered in 3D7 DiCre expressing *P. falciparum* parasites using a two-step CRISPR/Cas9-mediated gene editing approach (Supplementary Figure 2A). In the first step, a marker-free *loxP* site was introduced 6 base pairs upstream of the *myoa* start codon in the 5’ untranslated region (UTR) of the gene. In the second CRISPR/Cas9-mediated gene editing step, an mCherry tag was introduced at the 3’ end of the *myoa* gene, followed by a second *loxP* site and *hDHFR* cassette to select for integration. PCR analysis of the resulting Myo-cKO line confirmed the absence of wild type locus and correct gene editing at both editing sites within the gene (Supplementary Figure 2B).

The construct was designed so that addition of rapamycin (RAP) to MyoA-cKO parasites would lead to DiCre-mediated excision of the entire *myoa* open reading frame (Figure 2A). To examine the efficiency of excision, early ring stage parasites were treated for 4 hours with either 100 nM RAP or the equivalent volume of vehicle only (DMSO) and genomic DNA extracted from the resulting schizonts. PCR analysis confirmed that RAP treatment led to excision of the *myoa* gene, however a faint integration band could still be detected in the RAP-treated sample, indicating that excision was not 100% effective (Figure 2B). A faint excision band was also detected in the DMSO-treated sample, suggesting some stochastic activation of the Cre recombinase in the absence of RAP. Western blot analysis of these samples revealed a significant reduction in MyoA-mCherry signal at the protein level (Figure 2C), while immunofluorescence imaging confirmed complete loss of mCherry signal in 78.7% of RAP-treated parasites (Figure 2D), confirming that RAP-mediated excision was effective but incomplete across the parasite population. These findings are consistent with another published MyoA DiCre cKO line, which also reported incomplete excision following RAP treatment (Robert-Paganin et al., 2019).

**Figure 2.**
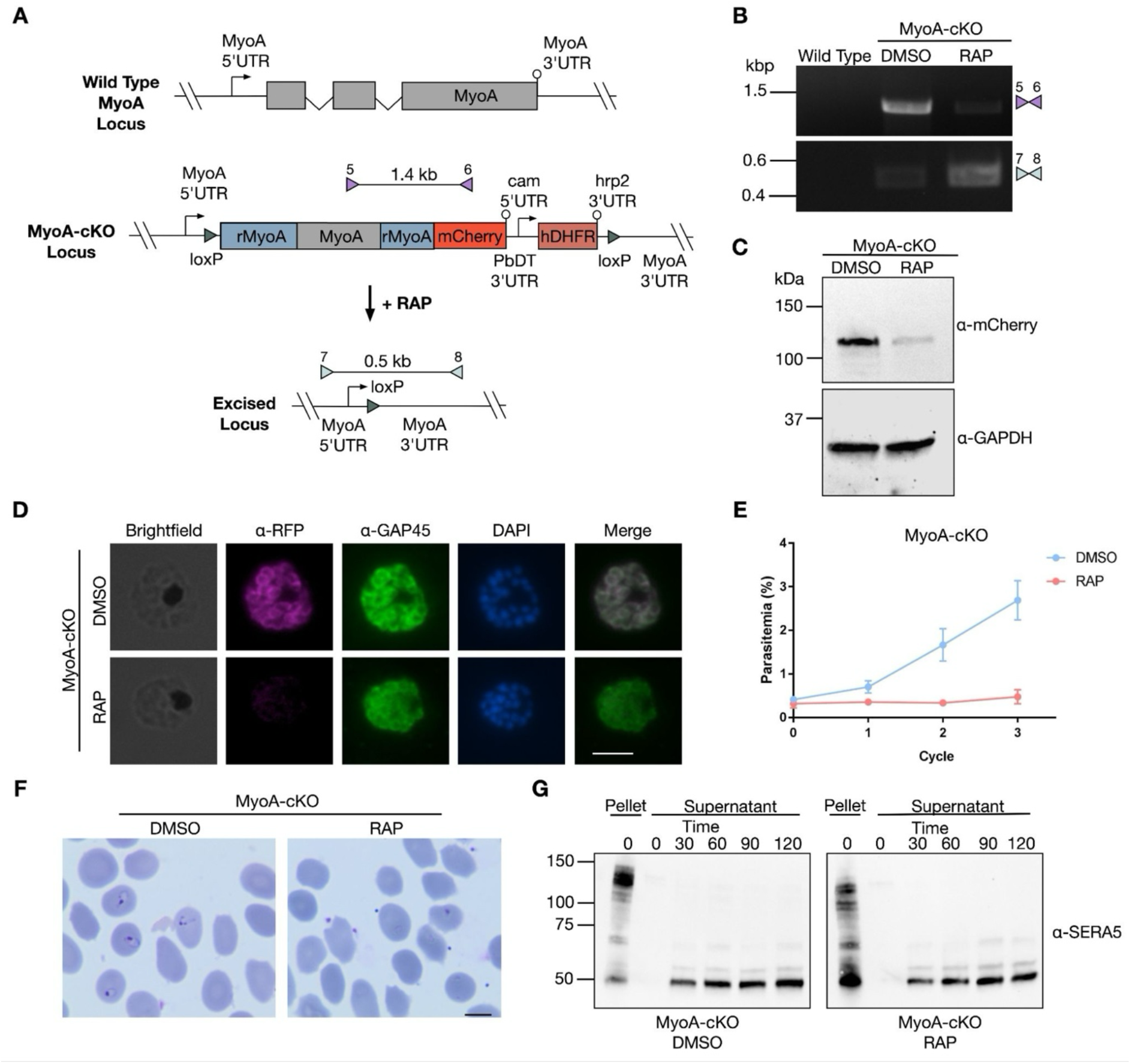
MyoA is essential for asexual parasite proliferation and is required for RBC invasion, but not egress. **A** Schematic of rapamycin-induced excision of the *myoA locus* in the MyoA-cKO line. Coloured triangles indicate primer pairs used for diagnostic PCR screens together with the expected amplicon sizes. **B** Diagnostic PCR analysis of WT and DMSO/RAP-treated MyoA-cKO parasites using the primer pairs shown in Fig.2A, assessing the unexcised locus (purple triangles) and the excised locus (teal triangles). **C** Western blot analysis of DMSO- and RAP-treated MyoA-cKO parasites probed with an anti-mCherry antibody. Blots were also probed with an anti-GAPDH as a loading control. **D** IFAs of DMSO- and RAP-treated MyoA-cKO parasites stained with anti-RFP and anti-GAP45 antibodies. Scale bar, 5 μm. **E** Growth curves of DMSO- and RAP-treated MyoA-cKO parasites over 3 intraerythrocytic growth cycles. Parasitemia was determined by flow cytometry. Data points plotted are means from two independent experiments, each performed in triplicate. Error bars represent the standard deviation. **F** Representative images of Giemsa-stained parasites from DMSO- and RAP-treated MyoA-cKO parasite cultures. Scale bar, 5 μm. **G** Western blot analysis monitoring the release of SERA5 into the culture supernatant of DMSO- and RAP-treated MyoA-cKO schizonts, as a measurement of egress over time. Pellet samples were included as loading controls.

To assess the impact of MyoA deletion on parasite growth, a flow cytometry-based growth assay was performed on DMSO- and RAP-treated MyoA-cKO parasites to monitor replication over three erythrocytic growth cycles. Consistent with previous reports (Blake et al., 2020; Robert-Paganin et al., 2019), MyoA-deficient parasites did not proliferate in culture, confirming that MyoA is essential for parasite growth (Figure 2E). Examination of Giemsa-stained blood smears from MyoA-cKO cultures 44 hours post excision showed an accumulation of free merozoites and an absence of ring stage parasites in RAP-treated cultures, demonstrating a defect in RBC invasion, but not in egress (Figure 2F). To confirm that loss of MyoA had no impact on egress dynamics, we monitored the appearance of proteolytically processed forms of the abundant parasitophorous vacuole protein serine repeat antigen 5 (SERA5) over time in culture supernatants from rupturing DMSO- and RAP-treated schizonts. Indeed, we observed no changes in the shedding kinetics of SERA5 (Figure 2G), confirming that deletion of MyoA does not affect merozoite egress, consistent with other studies that observed unimpaired egress following disruption of the glideosome (He et al., 2023; Moussaoui et al., 2020; Perrin et al., 2018; Robert-Paganin et al., 2019).

### MyoA is not required for glideosome assembly or gametocytogenesis

A previous study demonstrated that deletion of GAP45 led to a significant reduction in MTIP and MyoA levels, indicating that their stability is reliant on expression of GAP45 in *P. falciparum* (Perrin et al., 2018). This may serve as a mechanism to degrade excess MyoA and ensure that only glideosome-bound MyoA can bind to actin filaments. Having observed that GAP45 localisation and abundance was unaffected following deletion of MyoA (Figure 2D), we next investigated the effect of MyoA disruption on other glideosome components, including GAP50 and MTIP. IFA (Figure 3A) and western blot analysis (Figure 3B) on DMSO- and RAP-treated MyoA-cKO parasites revealed no changes in the localisation or abundance of these proteins following MyoA depletion, indicating that MyoA is dispensable for glideosome assembly and stability.

**Figure 3.**
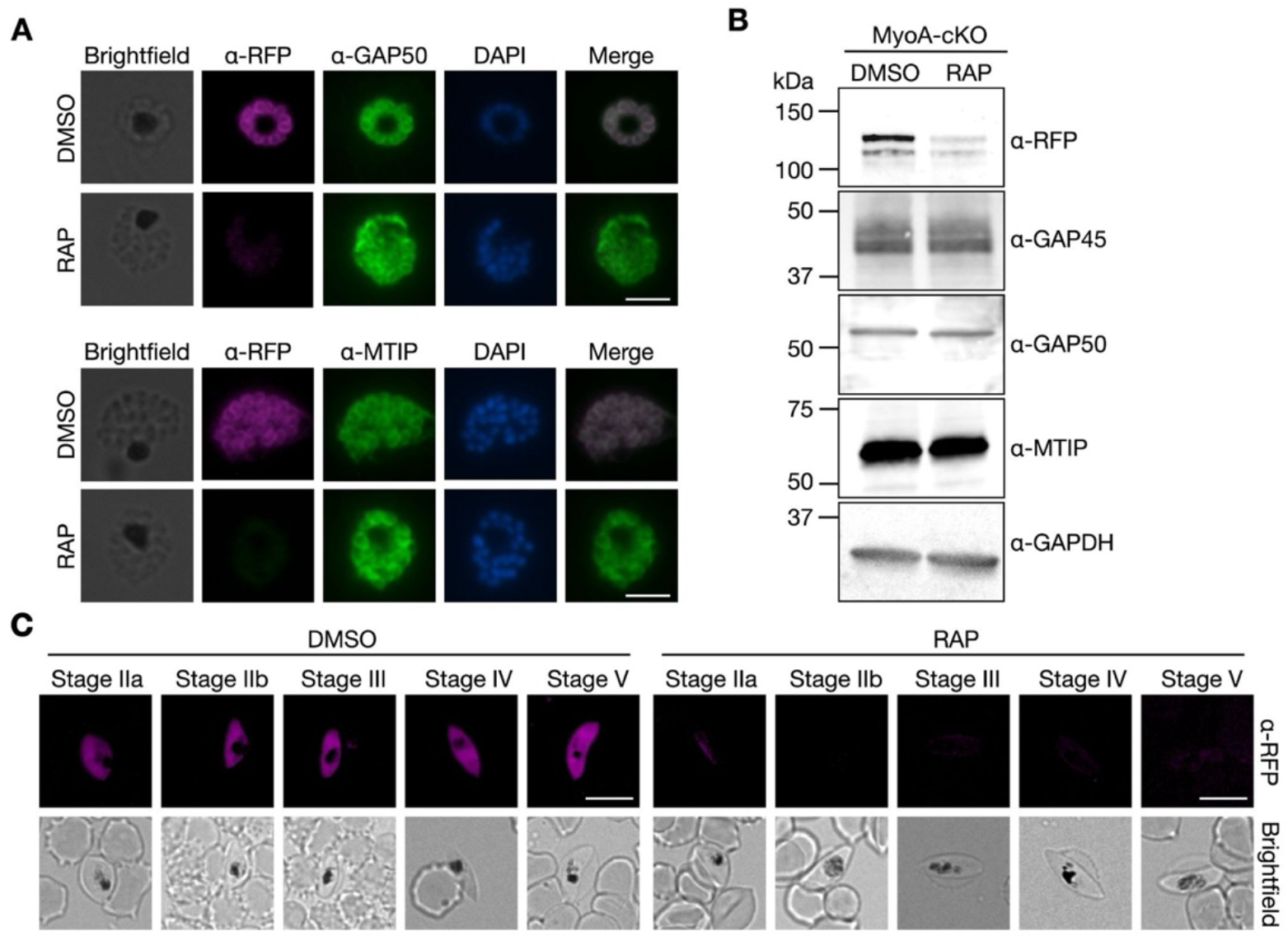
Loss of MyoA does not disrupt the expression levels of other glideosome components or gametocytogenesis. **A** IFAs of DMSO- and RAP-treated MyoA-cKO asexual parasites stained with antibodies against mCherry, GAP45, GAP50 and MTIP. Scale bars, 5 μm. **B** Western blot analysis of MyoA-mCherry, GAP45, GAP50, MTIP in DMSO- and RAP-treated MyoA-cKO parasites. Blots were also probed with anti-GAPDH as a loading control. **C** IFAs of DMSO- and RAP-treated parasites collected at the indicated stages of gametocyte development and stained with an anti-mCherry antibody. Scale bars, 10 μm.

While the role of the glideosome in generating motility during the asexual blood stages is well established, its function in gametocytes, the sexual stages of the lifecycle, remains unclear.

Gametocytes express core components of the glideosome and possess an IMC, which first forms as 13 disk-like structures that associate with the microtubules and progressively extend across the parasite as the gametocyte develops (Dearnley et al., 2011). Disruption of the IMC by knocking down proteins including PhIL1 and PIP1, leads to significant structural abnormalities in gametocytogenesis, with parasites unable to elongate (Parkyn Schneider et al., 2017). Similarly, knock down of GAP40, a structural component of the glideosome, leads to a reduction in gametocyte numbers and abnormal stage IV parasites which are unable to progress to stage V, highlighting an essential role for the glideosome in sexual development (He et al., 2023). While the IMC and the glideosome may provide structural support to the developing gametocytes, it remains unclear why the MyoA motor is expressed during this non-motile lifecycle stage (Dearnley et al., 2011).

To test if MyoA is also required for gametocytogenesis, we added RAP to sexually committed MyoA-cKO schizonts on day 3 of gametocyte induction, then monitored their development over two weeks. Live fluorescence microscopy revealed that MyoA is expressed at all stages of gametocyte development (Figure 3C). In RAP-treated parasites, some residual mCherry signal was still detectable in early stage IIa gametocytes, likely reflecting the late timing of RAP addition. In later stages, mCherry signal was no longer detectable and MyoA-deficient parasites were able to progress through gametocytogenesis and form mature stage V gametocytes without any discernible difference in morphology or overall gametocyte numbers when compared to DMSO-treated controls. Therefore, unlike GAP40, MyoA does not appear to be required for gametocyte development, suggesting that the glideosome fulfils a structural, rather than a motility-related function during this stage of the life cycle.

### Conditional complementation with wild type MyoA rescues the growth defect but S19A and S19D mutants rescue only partially

Having demonstrated that we could efficiently delete MyoA in our cKO line, we next wanted to complement the line with MyoA Ser19 mutants to investigate the role of this phosphosite on parasite growth. To ensure that the complemented copy of MyoA would only be expressed following excision of the endogenous gene, we adapted an inducible complementation approach (Patel et al., 2019). Compared to previous work, where either the endogenous gene was mutated (Ripp et al., 2022) or where a conditional MyoA cKO line was complemented with constitutively expressed ectopic MyoA mutants (Blake et al., 2020); our conditional system allows for temporal control, cleaner genetic backgrounds and reduces the likelihood of parasites adapting to the MyoA mutations over multiple cycles.

Briefly, we introduced an expression cassette consisting of a *myoa* promoter followed by a floxed sequence encoding GFP, a T2A viral skip peptide and the blasticidin-S deaminase (BSD) resistance gene into the *p230p* locus of our MyoA-cKO line. The floxed GFP sequence was then followed by a promoterless sequence encoding GFP-tagged WT MyoA or MyoA harbouring the S19A or S19D mutations giving rise to MyoA-cKO:compWT, MyoA-cKO:compS19A and MyoA-cKO:compS19D lines, respectively which would ensure that the mutant copies would not be expressed under normal culture conditions. RAP treatment of the three resulting parasite lines would lead to simultaneous excision of the endogenous mCherry-tagged MyoA and the floxed GFP sequence in the *p230p* locus, resulting in the deletion of the endogenous *myoa* gene and placement of the GFP-tagged complemented *myoa* sequence under the control of the *myoA* promoter (Figure 4A). PCR analysis of the resulting complemented lines confirmed integration of the complementation cassettes into the *p230p* locus and demonstrated the efficient excision of the floxed sequences in the endogenous *myoa* and *p230p* loci following RAP treatment (Figure 4B).

**Figure 4.**
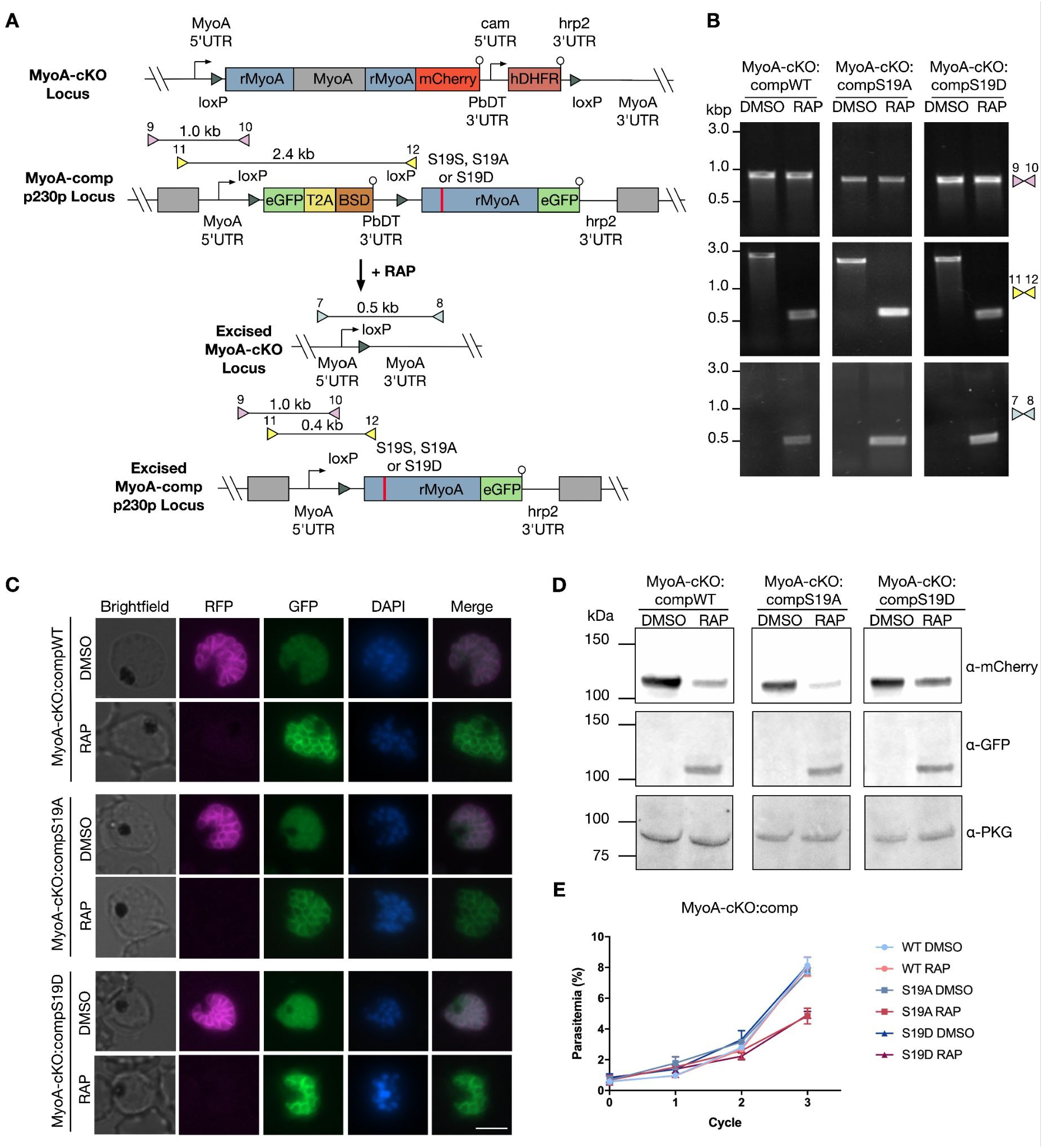
Complementation of MyoA-cKO parasites with Ser19 mutants of MyoA leads to defective parasite proliferation. **A** Schematic of the CRISPR/Cas9-mediated strategy used to generate conditional MyoA complementation lines in the MyoA-cKO background. A floxed GFP-T2A-BSDr cassette under the control of the *myoA* promoter was introduced into the *p230p* locus. This was followed by a promoterless sequence encoding GFP-tagged WT or mutant MyoA harbouring the S19A or S19D mutations, resulting in the generation of the MyoA-cKO:compWT, MyoA-cKO:compS19A and MyoA-cKO:compS19D lines. Rapamycin treatment induces excision of the endogenous MyoA-mCherry sequence as well as the GFP-T2A-BSDr sequence, placing the complemented copy of MyoA-GFP under the control of the *myoA* promoter in the *p230p* locus. Scissors indicate CRISPR/Cas9 cleavage sites. Coloured triangles indicate primer pairs used for diagnostic PCR together with the expected amplicon size. **B** Diagnostic PCR analysis of the MyoA-cKO:compWT, MyoA-cKO:compS19A and MyoA-cKO:compS19D lines using the primer pairs shown in Fig.4A, assessing 5’ integration of the complementation constructs at the *p230p* locus (pink triangles), excision of the GFP-T2A-BSDr cassette (yellow triangles) and excision of the endogenous MyoA-mCherry sequence (teal triangles). **C** Live fluorescence microscopy of DMSO- and RAP-treated MyoA-cKO:compWT, MyoA-cKO:compS19A and MyoA-cKO:compS19D parasites. **D** Western blot analysis of DMSO- and RAP-treated MyoA-cKO:compWT, MyoA-cKO:compS19A and MyoA-cKO:compS19D parasites probed with anti-mCherry and anti-GFP antibodies. Blots were also probed with anti-PKG as a loading control. **E** Growth curves of DMSO- and RAP-treated MyoA-cKO:compWT, MyoA-cKO:compS19A and MyoA-cKO:compS19D parasites. Parasitemia was determined by flow cytometry. Data are presented as the means from two independent experiments, each performed in triplicate. Error bars represent the standard deviation

To assess the efficiency of both endogenous mCherry-tagged MyoA deletion and activation of the GFP-tagged complementing MyoA, parasites were treated at the early ring stage with 100 nM RAP or the equivalent volume of DMSO for 3 hours and analysed by live fluorescence microscopy 44 hours later within the same cycle, when the parasites would be segmented schizonts. RAP-treated parasites exhibited a loss of mCherry signal, confirming deletion of the endogenous *myoa* gene. Simultaneously, recombination at the *p230p* complementation locus led to excision of the floxed GFP sequence, bringing the ectopic MyoA-GFP, MyoA(S19A)-GFP or MyoA(S19D)-GFP sequence under the control of the *myoa* promoter. As a result, the GFP signal switched from a cytosolic to a peripheral localisation following RAP treatment, consistent with the expected localisation of MyoA at the IMC (Figure 4C). Western blot analysis also confirmed that RAP treatment resulted in induction of MyoA-GFP expression as well as a reduction, but not total loss of MyoA-mCherry expression (Figure 4D). Although GFP signal was detectable by microscopy in the DMSO-treated parasites (Figure 4C), the expected ∼42 kDa GFP band was not detectable by western blot(Figure 4D), likely reflecting the low abundance of the cytosolic GFP signal in DMSO-treated parasites.

A flow cytometry-based growth assay demonstrated that RAP-treated MyoA-cKO:compWT parasites replicated at the same rate as control DMSO-treated parasites (Figure 4E), confirming efficient genetic complementation of the endogenous *myoa* gene deletion. In contrast, RAP-treated MyoA-cKO:compS19A parasites showed a 37% reduction in growth following 3 replication cycles (Figure 4E). This is in line with a previous study that demonstrated a 55% reduction in parasite growth when MyoAcKO parasites were complemented with a MyoA mutant harbouring a K764E mutation that blocks the electrostatic interaction with the phosphorylated Ser19 residue (Blake et al., 2020). Surprisingly, we also observed a 40% reduction in the growth of RAP-treated MyoA-cKO:compS19D parasites (Figure 4E), despite previously showing that the S19D mutation was tolerated in the endogenous *myoa* gene (Figure 1F). These findings suggest that constitutively phosphorylated MyoA could be deleterious to parasite growth, leading to the observed growth defect in RAP-treated MyoA-cKO:compS19D parasites. However, it is also possible that the S19D mutation does not functionally mimic phosphorylated Ser19 and therefore behaves like a phosphomutant instead.

### *In vitro* motility assays reveal that S19A reduces the gliding speed of the motor

Having demonstrated that the MyoA S19A and S19D mutations resulted in defective parasite growth, we next wanted to test whether the phosphorylation status of MyoA and these mutations affected the activity of the motor. To do so, we performed *in vitro* motility assays using purified MyoA-mCherry that was episomally expressed in parasites (Figure 5A). In these assays, we tracked the motility of ATP-driven movement of rabbit actin filaments labelled with rhodamine-phalloidin over glass cover slips after coating them with mCherry-tagged MyoA captured from mature schizont or merozoite lysates with an anti-mCherry antibody (Figure 5B). Motility assays were performed using MyoA-mCherry derived from C2- and E64-blocked schizonts and purified merozoites to test whether blocking PKG-dependent phosphorylation of MyoA Ser19 had an effect on filament motility. We found that the average filament velocity for C2-treated samples was 0.52 (±0.13) μm/s, while E64-treated samples moved faster with an average velocity of 0.55 (±0.14) μm/s (Figure 5C). Merozoite samples, which had the highest phosphorylation levels of MyoA Ser19, displayed an average filament velocity of 0.60 (±0.18) μm/s; 14% and 8% faster than C2- and E64-blocked samples, respectively. Therefore, there appears to be a correlation between motor speed and phosphorylation levels of MyoA Ser19 since merozoite samples display the highest levels of Ser19 phosphorylation and actin displacement velocities, followed by E64 then C2-blocked samples. However, since MyoA co-immunoprecipitates with other components of the glideosome, including GAP40 and GAP45 (Green et al., 2017), which are also phosphorylated in a PKG-dependent manner (Alam et al., 2015), it remained possible that the changes in actin filament velocity could be due to the altered phospho-status of other glideosome components and not solely due to the phospho-status of MyoA Ser19.

**Figure 5.**
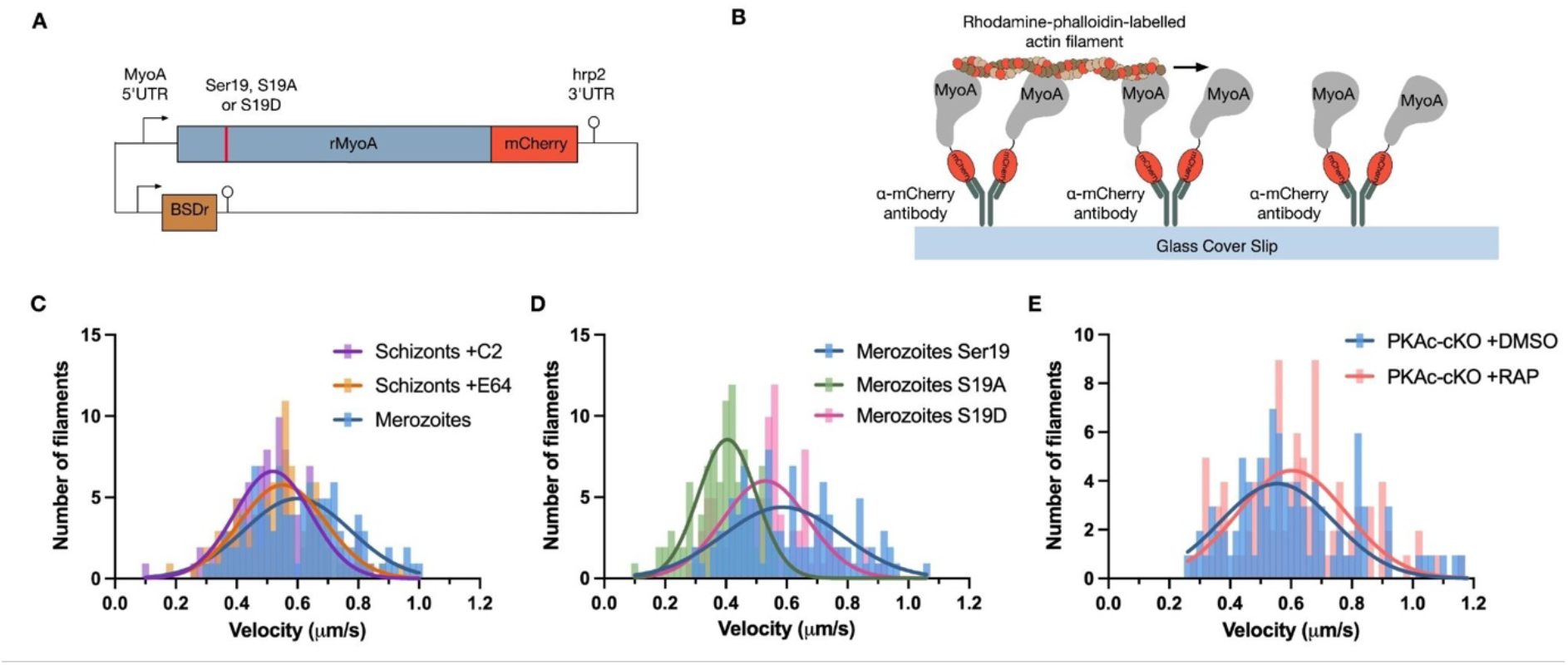
Phosphorylation of MyoA enhances actin motility in *in vitro* gliding assays. **A** Schematic of the plasmid used for episomal expression of mCherry-tagged MyoA, including wild type, phosphomutant (S19A) and phosphomimetic (S19D) variants. **B** Schematic of the *in vitro* motility assay flow cell set up, showing rhodamine-phalloidin labelled actin moving over parasite-derived MyoA-mCherry captured via an anti-mCherry antibody on glass cover slips. **C** Speed distributions from *in vitro* motility assays using MyoA-mCherry isolated from C2-arrested schizonts(N=102), E64-arrested schizonts (N=105) or merozoites (N=102). Data are derived from three independent biological sample preparations. **D** Speed distributions from *in vitro* motility assays using MyoA(S19)-mCherry (N=104), MyoA(S19A)-mCherry (N=104) or MyoA(S19D)-mCherry (N=105) derived from merozoites. Data are from three independent biological sample preparations. **E** Speed distributions from *in vitro* motility assays using MyoA(S19)-mCherry derived from PKAc-cKO merozoites treated with DMSO (N=91) or RAP (N=101). Data are from three independent biological sample preparations.

To address this issue and to ensure that the only difference between samples was the phospho-status of MyoA, wild type 3D7 parasites were transfected with plasmids driving episomal expression of MyoA(Ser19)-mCherry, MyoA(S19A)-mCherry or MyoA(S19D)-mCherry, while the endogenous MyoA locus was left unaltered (Figure 5A). The *in vitro* motility assays were performed using MyoA-mCherry immunoprecipitated from merozoites to ensure maximal S19 phosphorylation levels in the control wild type samples. The average filament velocity for wild type MyoA(Ser19)-mCherry samples was 0.59 (±0.20) μm/s, while the phosphomutant MyoA(S19A)-mCherry samples moved 31% slower, with an average velocity of 0.41 (±0.10) μm/s (Figure 5D), similar to that of non-phosphorylated MyoA from C2-treated schizonts in the assays performed above (Figure 5C). The phosphomimetic variant, MyoA(S19D)-mCherry, displayed an intermediate velocity of 0.53 (±0.14) μm/s, indicating that this mutation partially restores motor activity to that of the wild type phosphorylated MyoA (Figure 5D).

Finally, we wanted to test whether perturbations to cAMP signalling also affect MyoA motor activity, given that conditional disruption of PKAc results in an invasion defect resembling that of the MyoA knockout (Patel et al., 2019). PKAc was initially proposed to directly phosphorylate MyoA at Ser19 (Lasonder et al., 2012); however, this model was challenged by the finding that the MyoA phosphosite is still detected in PKAc knockout parasites (Patel et al., 2019), suggesting that another kinase that acts downstream of PKG, such as a calcium-dependent protein kinase (CDPK) is instead responsible for phosphorylating Ser19. However, elevated cAMP levels in PDEβ knockout parasites promote MyoA phosphorylation in a PKG-independent manner, despite no detectable increase in cytosolic calcium levels (Flueck et al., 2019). Together with the observation that the PKA inhibitor H89 reduces MyoA phosphorylation (Flueck et al., 2019), these findings raise the possibility that PKA signalling may contribute to the regulation of MyoA phosphorylation under certain conditions. To investigate this, we introduced the episomal MyoA(Ser19)-mCherry construct (Figure 5A) into the PKAc conditional knockout line generated previously (Patel et al., 2019). Parasites were treated at the ring stage with either DMSO to maintain PKAc expression, or with RAP to induce PKAc deletion. Merozoites were subsequently purified and used for *in vitro* motility assays as described above. The average filament velocity for DMSO-treated parasites was 0.62 (±0.20) and 0.63 (±0.19) μm/s for RAP-treated PKAc knockout parasites (Figure 5E, indicating that loss of PKAc does not measurably alter MyoA motor speed under the conditions tested.

## Discussion

In this study, we demonstrate that MyoA is essential for asexual parasite growth but is dispensable for gametocyte development. We further show that dynamic phosphorylation of MyoA at Serine 19 (Ser19) is a crucial regulatory mechanism governing the speed of the *P. falciparum* motor and is also critical for optimal parasite proliferation during the asexual blood stages of infection. Building on previous phosphoproteomic analyses that identified Ser19 as a target of PKG-dependent phosphorylation (Alam et al., 2015; Flueck et al., 2019), we show that Ser19 phosphorylation occurs rapidly and transiently, peaking in extracellular merozoites and decreasing dramatically in ring stages post-invasion. This temporal regulation suggests that MyoA phosphorylation is tightly coupled to parasite egress and invasion and must be precisely regulated for optimal motor function.

Using an inducible knockout system complemented with conditionally expressed wild type or phosphomutant MyoA, we show that dysregulation of Ser19 phosphorylation, either by blocking phosphorylation (S19A) or mimicking constitutive phosphorylation (S19D), impairs parasite growth. While S19A complementation partially rescues the growth defect, the parasites’ reduced replication rate and the diminished *in vitro* motor activity of MyoA(S19A) reinforce the importance of phosphorylation at this residue for MyoA function. Intriguingly, despite the phosphomimetic S19D mutation restoring *in vitro* motor speed to near wild type levels, it also results in a significant growth defect, similar to that of the S19A mutant, suggesting that timely dephosphorylation of MyoA is equally critical for successful invasion and intracellular development. This observation raises the possibility that S19D, although capable of maintaining electrostatic interactions with the Lys764 residue (Robert-Paganin et al., 2019), may lock the motor in a constitutively active state that disrupts temporal coordination with other invasion processes. However, phosphomimetic mutations are not always successful at reconstituting phosphorylated residues, since the chemical environment created by negatively charged amino acids such as aspartic acid is different to the chemical environment of phospho-serine (Hunter, 2012). Therefore, it is also possible that the S19D mutation introduces subtle structural perturbations or fails to fully recapitulate the dynamic reversibility of phosphorylation, both of which could compromise glideosome function within the parasite.

Our conditional complementation approach demonstrates that only wild type MyoA fully rescues the growth defect of the MyoA knockout, emphasizing the importance of regulatory plasticity at Ser19. This aligns with previous work showing that perturbing the electrostatic interaction between phosphorylated Ser19 and Lys764 by introducing a K764E mutation reduces invasion efficiency and results in defective internalisation (Blake et al., 2020). Together with our findings, this highlights the role of this interaction as a molecular tuning switch for motor activity. Furthermore, our use of chemical inhibitors to modulate MyoA phosphorylation and gliding speed *in vitro* further support the role of Ser19 phosphorylation as a reversible mechanism used to regulate motor speed.

Importantly, this is the first study to assess MyoA phosphorylation mutants using parasite-derived MyoA rather than recombinant MyoA. The discrepancies in gliding motility speeds between our findings and previously published gliding assays likely reflect inherent biological differences between the simplified recombinant systems, which only consist of MyoA, ELC, and MTIP and the native MyoA complexes, which co-immunoprecipitate with more glideosome-associated proteins including GAP40, GAP45 and GAP50 (Baum et al., 2005; Green et al., 2017). Future studies should aim to dissect the contribution of additional glideosome components to motility regulation, particularly since several of these proteins are also phosphorylated in a PKG-dependent manner prior to egress (Alam et al., 2015). Interestingly, a recent study demonstrated that introduction of S370A, S372A and S376A mutations in PfGAP45 impairs merozoite invasion (He et al., 2023). Although these phosphosites are not PKG-dependent (Alam et al., 2015), these findings provide further evidence that phosphorylation-mediated regulation of the glideosome is critical for efficient parasite invasion.

Although our study clearly establishes the regulatory importance of Ser19 phosphorylation, the identity of the kinase that acts downstream of PKG to phosphorylate MyoA, as well as the phosphatase involved in dephosphorylating the motor post-invasion remain unknown. Identifying these proteins will be key to fully elucidating the signalling cascade controlling glideosome activation, deactivation and possibly disassembly. Although we found that depletion of PKA had no measurable effect on motor speed, these experiments demonstrate that the parasite-derived *in vitro* motility assay provides a robust platform for assessing the functional consequences of genetic perturbations on the glideosome. This approach should prove valuable for dissecting how additional invasion regulators influence glideosome function.

In conclusion, our data reveal that MyoA phosphorylation at Ser19 acts as a reversible switch that regulates motor activity and is essential for efficient parasite invasion and proliferation. The timing and reversibility of this phosphorylation are critical, with both hypo- and hyperphosphorylated states leading to impaired growth. This study provides new mechanistic insight into glideosome regulation and highlights the potential of targeting cyclic nucleotide signalling and MyoA phosphoregulation as antimalarial strategies.

## Methods

### *In vitro* culture and synchronisation of *P. falciparum* parasites

*P. falciparum* 3D7a DiCre asexual blood stages were cultured in human erythrocytes of various blood groups (National Blood Transfusion Service, London, United Kingdom) using complete medium consisting of RPMI-1640 medium (Life Technologies) supplemented with 0.5% AlbuMAX type II (Gibco), 50 μM hypoxanthine and 2 mM L-glutamine. Parasite cultures were incubated at 37°C and gassed with 90% N_2_, 5% CO_2_ and 5% O_2_ according to standard procedures (Trager & Jensen, 1976). Parasitaemias were routinely monitored by light microscopy examination of thin blood films fixed with 100% methanol and stained with 10% Giemsa stain in phosphate buffer (8 mM KH_2_PO_4_, 6 mM Na_2_HPO_4_, pH 7.0).

Tightly synchronous parasites were obtained by purifying segmented schizonts on a cushion of 70% Percoll (GE Healthcare) and allowing them to invade fresh erythrocytes for 1-2 hours in a shaking incubator. This was followed by lysis of unruptured schizonts by treating with 5% D-sorbitol (Sigma) for 10 minutes at 37°C (Lambros & Vanderberg, 1979) to obtain highly pure and synchronous ring stage cultures.

### Gametocyte induction and culture

Induction of gametocytes was achieved using an adapted version of previously described techniques (Fivelman et al., 2007). Briefly, on Day 0, highly synchronous ring-stage parasites at 8-10% parasitemia were subjected to nutrient stress by replacing only half of the spent culture medium with fresh medium. On Day 1, half the medium was again replaced with fresh culture medium. From Day 2 onwards, parasite cultures were fed daily with fresh pre-warmed medium. To prevent asexuals parasite replication, heparin was added to the cultures at a final concentration of 20 U/ml from Day 3 onwards.

### Parasite transfection

Highly synchronous late-stage schizonts were transfected as previously described (Collins et al., 2013) using the AmaxaTM 4D-Nucleofector system (Lonza) and program FP158. For each transfection 20-50 μg plasmid DNA was resuspended in 100 μL of supplemented P3 primary cell solution, which was then used to resuspend 25 μL of Percoll purified schizonts (∼1.25x10^8^ cells) obtained from synchronous cultures. Parasites were electroporated using the FP158 setting then transferred back into culture and placed on the shaker for 1-2 hours. Appropriate drug selection was applied 24 hours post transfection. To select for parasites harbouring either the hDHFR, BSDr or yDHODH selection cassettes, 2.5 nM WR (Jacobus Pharmaceuticals), 2.5 μg/ml BSD (Sigma-Aldrich) or 1.5 μM DSM1 (MR4/BEI Resources) were used, respectively. For marker-free CRISPR/Cas9 transfections, parasites were treated with the appropriate drug based on the selection cassette found on the Cas9 plasmid for 6 days. For episomal transfections or CRISPR/Cas9 transfections where a drug selection cassette was integrated into the genome, parasites were maintained under constant drug selection pressure.

Once parasites were observed by Giemsa smear and integration confirmed by PCR (see Supplementary Table 1 for all primers), clonal parasite lines were obtained by limiting dilution and assessing for single plaque formation as previously described (Thomas et al., 2016).

### Plasmid construction

Benchling’s CRISPR gRNA design software (www.benchling.com), was used to identify suitable CRISPR/Cas9 targeting sites in the *P. falciparum* genome based on the sequences deposited in PlasmoDB for Myosin A (PF3D7_1342600) and P230p (PF3D7_0208900). gRNA sequences were selected based on lowest probability of off-target cleavage and closest proximity to the homology regions used to repair the double stranded break. A list of gRNAs used in this study are available in Supplementary Table 1. gRNA CRISPR/Cas9 plasmids were generated by introducing the desired gRNA sequences into either the pDC2-Cas9 harbouring the hDHFR cassette (Knuepfer et al., 2017) or the pL6 plasmid (Ghorbal et al., 2014).

The repair plasmid to generate the loxP:MyoA line was generated by amplifying the 5’ and 3’ homology regions (both 1 kb) from *P. falciparum* genomic DNA, while the sequence encoding a loxP site and ∼250 bp of recodonised *MyoA* sequence was ordered from Integrated DNA technologies (IDT). The three fragments were InFusion cloned into a pUC19 vector. This plasmid was further modified by inverse PCR to introduce the S19A and S19D mutations.

To generate the endogenously tagged MyoA-mCherry and MyoA-cKO lines, a repair plasmid was made by amplifying the 5’ (1 kb) from *P. falciparum* genomic DNA, and InFusion cloned into the pL6 plasmid(Ghorbal et al., 2014), along with a ∼400 bp of recodonised *MyoA-mCherry* sequence ordered from Integrated DNA technologies (IDT). A subsequent InFusion cloning step introduced the *hDHFR* expression cassette, followed by a *loxP* site and the 3’HR (1 kb). In a final cloning step, the sgRNA was introduced by ligation-based cloning.

To generate the conditional complementation repair constructs in the *P230p* locus, the plasmid described by Patel et al., was modified to replace the *cam* promoter with the *myoA* promoter, which comprised ∼2700 bp upstream of the *myoA* coding sequence. The plasmid was also further modified to replace the PKAc-3xHA-DDD sequence with recodonised MyoA-GFP sequence ordered from IDT. This plasmid was further modified by inverse PCR to introduce the S19A and S19D mutations.

Episomal expression constructs for the *in vitro* motility assays were made by introducing the *myoA* promoter, which comprised ∼2700 bp upstream of the *myoA* coding sequence, followed by recodonised MyoA-mCherry into a pUC19 plasmid by InFusion cloning. This plasmid was further modified by inverse PCR to introduce the S19A and S19D mutations.

### Western blot analysis

Parasites were released from erythrocytes by lysing in 5 volumes of 0.15% saponin (Sigma) in PBS containing cOmplete EDTA-free protease inhibitor (Roche) and 1 mM PMSF (Thermo Fisher Scientific). The samples were pelleted at 12,000 x g for 1 minute then washed twice in PBS also containing cOmplete EDTA-free protease inhibitor (Roche) and 1 mM PMSF (Thermo Fisher Scientific). The sample pellets were snap frozen in a dry ice/ethanol slurry and stored at -80°C. Saponin-released parasite pellets were lysed in 4 volumes of CoIP buffer containing 150 mM NaCl, 10 mM Tris (pH 7.5), 0.5 mM EDTA and 1% NP40 that was supplemented with cOmplete EDTA-free protease inhibitor (Roche) and 1 mM PMSF (Thermo Fisher Scientific) and incubated on ice for 10 minutes. Samples were centrifuged at 12,000 x g for 10 minutes at 4°C and the supernatant collected. Reducing sample buffer was added to NP40-lysed parasite samples or culture supernatants and proteins were resolved on 4%-15% Mini-PROTEAN TGX Stain-Free Precast Gels (Bio-Rad). Proteins were transferred onto nitrocellulose membranes using a semidry Trans-Blot Turbo Transfer System (Bio-Rad) and blocked using 10% skimmed milk in PBS containing 0.1% Tween-20 (PBST) for 1 hour and subsequently probed with either rat anti-RFP (1:5,000; Clone 6G6; Chromotek), mouse anti-GAPDH (1:30,000; gifted by Claudia Daubenberger (Swiss Tropical and Public Health Institute, Switzerland)), rabbit anti-PKG (1:1,000; Enzo Life Sciences), rabbit anti-mCherry (1:2,500; ab183628; Abcam), rabbit anti-SERA5 (1:5,000, (Stallmach et al., 2015)), rabbit anti-MyoA, GAP45, GAP50 or MTIP (1:5,000; gifted by Judith Green at the Francis Crick Institute, UK (Ridzuan et al., 2012), rabbit anti-pSer19 MyoA (1:1,000; raised against the phosphopeptide ‘N’-RRV[pS]NVEAFDKC (Alam et al., 2015)). After probing with primary antibodies for at least 1 hour, membranes were washed three times for 5 minutes in PBST followed by incubation with the relevant secondary antibodies conjugated to near infrared (NIR) dyes for 1 hour (1:5,000; Invitrogen Molecular Probes). Membranes were washed three times for 5 minutes in PBST followed by a single wash in PBS. Membranes were dried between Whatman 3MM blotting papers and images using an Azure c600 Imaging System (Azure Biosystems) or a ChemiDoc Imaging System (Bio-Rad). Densitometry quantifications were performed using ImageJ and plotted in GraphPad Prism.

### Live parasite imaging

To image live parasites, 1 ml of parasite culture was stained with 1 μg/ml Hoechst 33342 for 5 minutes at 37°C to visualise nuclei. The parasites were then washed three times in RPMI, and a small amount of culture material at ∼50% haematocrit was added to a microscope slide and covered with a coverslip. Images were acquired using an EVOS FL cell imaging system and processed using ImageJ.

### Microscopy of Giemsa-stained blood films

Thin blood films of parasite cultures were fixed using 100% methanol and stained for 5 minutes with 10% Giemsa stain (Merck). Images were obtained using an Olympus BX51 microscope fitted with an Olympus SC30 digital colour camera through a 100X oil immersion objective.

### Immunofluorescence analysis (IFA)

Thin blood smears were fixed using high-quality methanol-free 4% formaldehyde (Thermo Scientific) in PBS for 20 minutes, followed by two PBS washes then permeabilised with 0.1% Triton X-100 in PBS for 10 minutes. Detergent was washed off twice in PBS and slides were blocked in 3% bovine serum albumin (BSA) in PBS for 1 hour and subsequently probed for 1 hour with either rat anti-RFP (1:1,000; Clone 5F8; Chromotek), rabbit anti-mCherry (1:500; ab183628; Abcam), rabbit anti-GAP45, GAP50 or MTIP (1:1,000; gifted by Judith Green (Ridzuan et al., 2012)). Slides were washed three times with PBS then probed with the relevant AlexaFluor secondary antibodies (1:500; Invitrogen Molecular Probes) in blocking solution for 1 hour. Slides were washed three times before being mounted in ProLong™Gold Antifade Mountant containing DAPI (Thermo Fisher Scientific). Images were acquired using either a Nikon Eclipse Ti fluorescence microscope fitted with a Hamamatsu C11440 digital camera, or an EVOS® FL Cell Imaging System and processed using ImageJ.

### Flow cytometry-based growth assays

To measure parasite growth over time by flow cytometry, parasites were synchronised to a 4-hour window. Cultures were adjusted to 0.1-0.5% parasitaemia in 4% haematocrit in triplicate and treated either with 100 nM rapamycin or the equivalent volume of DMSO for 4 hours. Samples were taken every 48 hours and fixed using high-quality methanol-free 4% formaldehyde (Thermo Scientific), 0.1% glutaraldehyde (Sigma) in PBS for 1 hour. Fixed samples were stained with SYBR Green I (Molecular Probes) then analysed using a BD LSR II flow cytometer (BD Biosciences), with 50,000 events collected for each sample. FlowJo 7 analysis software (FlowJo LLC) was used to analyse the data. Different assays were adjusted for starting parasitaemias and growth curves were generated using GraphPad Prism.

### Egress timepoint assays

Parasites were synchronised to a 2-hour window and treated for 4 hours with either 100 nM RAP or the equivalent volume of DMSO. At ∼24 hpi at the early trophozoite stage, parasites were treated with 1.5 μM C2 overnight. At ∼48 hpi, once mature segmented schizonts were observed, the parasites were Percoll enriched and washed several times in pre-warmed RPMI. Parasites were resuspended in RPMI at 3.25x10^8^ parasites/ml and 65 μl aliquots were dispensed in five 1.5 ml eppendorf tubes. The parasites were then incubated at 37°C for 0, 30, 60, 90 or 120 minutes. To harvest samples at each time point, parasites were pelleted at 9,000 x g and culture supernatants were purified using 0.22 μm Costar Spin-X centrifuge filters (Corning) to remove free merozoites and contaminating parasite material. The parasite pellet from the first time point was retained as a parasite loading control. Samples were subject to western blot analysis and probed with a rabbit anti-SERA5 (1:5,000 (Stallmach et al., 2015)) antibody as a measure of schizont egress.

### Preparation and purification of merozoites

Schizonts from tightly synchronised cultures were purified by magnetic cell separation (MACS) enrichment (Mata-Cantero et al., 2014). Briefly, a D column (Miltenyi Biotech) was fitted with a two-way stopcock attached to a 22-gauge blunt-ended needle with a protective cover to avoid needle pricks. The column was then placed in a SuperMACS™II separator (Miltenyi Biotech) and washed with 10 ml of RPMI. Percoll-purified schizonts were resuspended in 4 ml of RPMI and added to the column and then washed twice with 4 ml RPMI. The column was then removed from the magnet, and schizonts eluted in 8 ml of RPMI. Schizonts were then left to rupture in a shaking incubator for 1 hour. To separate merozoites from unruptured schizonts, the cultures were then re-loaded onto the D column placed in a SuperMACS™ II separator which would retain unruptured schizonts and hemozoin-associated cell debris on the magnet and allowing merozoites to elute in the flowthrough. This was followed by a wash step with 4 ml of RPMI. The flowthrough was then centrifuged at 3,500 x g for 10 minutes and supernatants discarded. Merozoites samples were snap frozen in a dry ice/ethanol slurry and stored at -80°C until required.

### *In vitro* motility assays

Motility assays were performed as previously described (Butt et al., 2010). Motility assay buffer (AB) containing 25 mM imidazole-HCL, 25 mM KCl, 1 mM EGTA and 4 mM MgCl_2_ (pH7.4) was repeatedly degassed and flushed with nitrogen and stored in a hypodermic syringe to prevent oxygen contamination which could lead to photobleaching ((Green et al., 2006). Merozoite and saponin-released schizont pellets were lysed in 4 volumes of CoIP buffer containing 150 mM NaCl, 10 mM Tris (pH 7.5), 0.5 mM EDTA and 1% NP40 that was supplemented with cOmplete EDTA-free protease inhibitor (Roche) and 1 mM PMSF (Thermo Fisher Scientific). Samples were incubated on ice for 10 minutes then centrifuged at 12,000 x g for 10 minutes at 4°C and the supernatant collected and stored on ice while the motility assay flow-cell chambers were prepared.

Custom flow-cell chambers with an internal volume of ∼10 μl were constructed by adhering nitrocellulose-coated cover slips to microscope slides with strips of double-sided tape. Anti-mCherry antibody (ab183628, Abcam) was diluted 1:50 to 7 μg/ml in AB and was applied in the flow-cell for 5 minutes. Unbound antibody was washed out with AB then blocked for 5 minutes with AB containing 0.5 mg/ml BSA (AB/BSA). Parasite lysates were applied into the flow-cell and incubated for 5 minutes to capture MyoA-mCherry complexes. Unbound material was washed out twice with AB/BSA. AB containing 0.1 μM rhodamine-phalloidin-labelled F-actin was applied to the flow-cell and incubated for 2 minutes. Unbound actin filaments were washed out using AB/BSA. Imaging buffer (IB) was made using AB/BSA containing 20 mM DTT, 0.2 mg/ml glucose oxidase, 0.5 mg/ml catalase, 3 mg/ml glucose to scavenge oxygen and prevent photobleaching. The flow-cell was washed twice with IB then viewed by fluorescence microscopy to check for actin binding. This was done using an Axioskop 40 fluorescence microscope with a Zeiss PlanNeofluar x100, 1.3 numerical aperture objective lens. Fluorescence was excited by a mercury arc lamp using a rhodamine filter set (excitation filter HQ535/50, dichroic mirror Q565LP, and emission filter HQ605/75; Chroma Technology), and light emitted from the rhodamine-phalloidin-labelled actin filament specimen was imaged using an IC-310 image-intensified charge-coupled device camera (Photon Technology International). In order to activate the myosin motors, IB containing 2 mM ATP and 1% methylcellulose (IB/ATP) was applied to the flow-chamber. Experimental temperature was 23°C, Sequences of video frames were captured every 40 ms using a frame grabber card (Multipix Imaging Limited). Each image sequence was saved as a median of five consecutive image blocks, and filaments were manually tracked with GMimPro (Mashanov & Molloy, 2007). Non-motile or short tracks were removed from the analysis. From the resulting velocities, frequency distribution histograms were calculated (bin size of 0.02 μm/s), and a Gaussian curve was fitted with GraphPad Prism version 7.

## Acknowledgements

We thank members of the Baker, Blackman, Moon and Treeck labs for helpful discussions and critical input. We also thank Judith Green for providing the antibodies against MyoA, GAP45 & MTIP, as well as Claudia Daubenberger for the GAPDH antibody.

This work was supported by a joint Wellcome Trust Senior Investigator Award to D.A.B (106240/Z/14/Z) & M.J.B (106239/Z/14/A) and Wellcome ISSF2 funding to the London School of Hygiene and Tropical Medicine. This work was also supported by funding to M.J.B and J.E.M from the Francis Crick Institute, which receives core funding from CRUK, the MRC and the Wellcome Trust (FC001043 & FC001119). S.D.N was supported by BBSRC LIDo PhD studentship (BB/M009513/1).

## Author contributions

Conceptualisation: S.D.N, M.J.B, C.F, D.A.B

Formal analysis, Investigation, Visualisation, Methodology, Writing - original draft: S.D.N

Resources: J.E.M

Supervision: K.K, J.V, J.E.M, M.J.N, D.A.B

Funding Acquisition: S.D.N, M.J.B, D.A.B

Writing – review and editing: S.D.N., K.K, J.V, I.K, J.E.M, M.J.B, C.F, D.A.B

## Competing interests

The authors declare no competing interests

---

**Supplementary Figure 1.**
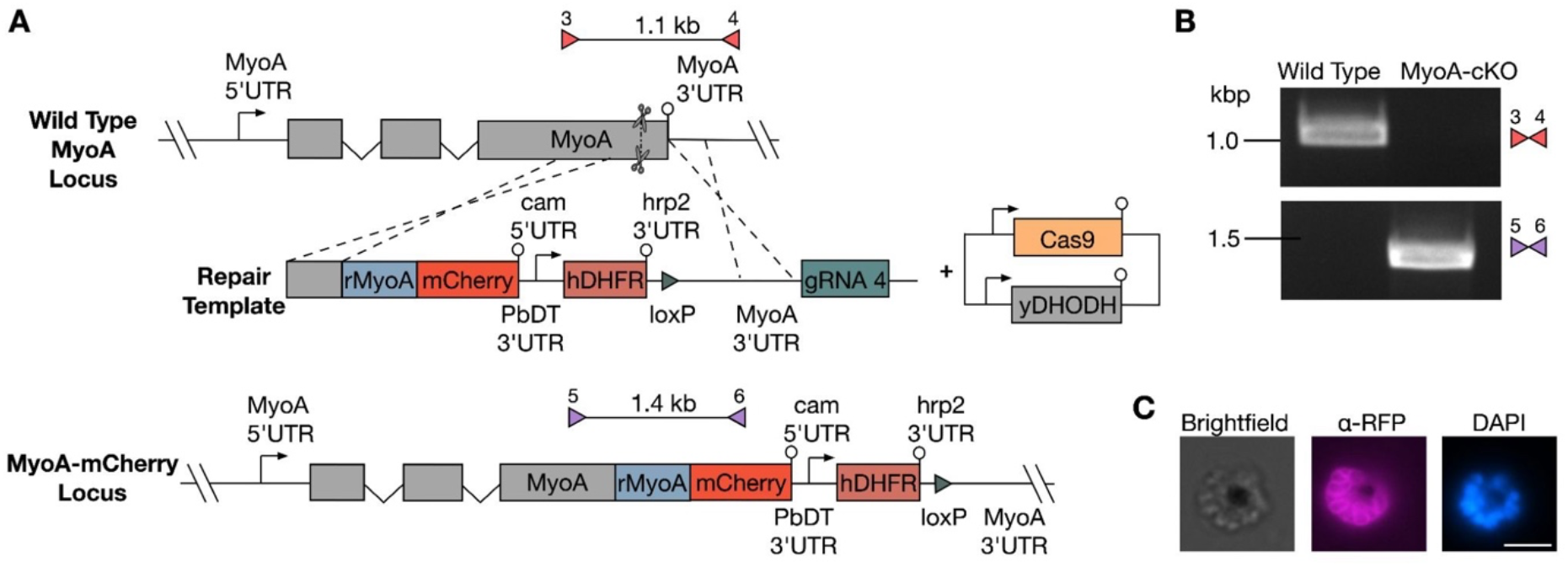
Generation of the MyoA-mCherry line. **A** Schematic of the CRISPR/Cas9-mediated approach used to generate the MyoA-mCherry line. A linear repair template introduces the C-terminal mCherry tag, a loxP site and an hDHFR cassette to select for integration. Scissors indicate CRISPR/Cas9 cleavage sites, while coloured triangles represent primer pairs used for diagnostic PCR screens, together with the expected amplicon sizes. **B** Diagnostic PCR analysis of the parental WT and MyoA-mCherry lines using the primer pairs shown in Supp.Fig1A, assessing loss of the WT locus (red triangles) and 5’ integration of the mCherry tag (purple triangles). **C** IFAs of MyoA-mCherry parasites stained with an anti-RFP antibody. Scale bar, 5 μm.

**Supplementary Figure 2.**
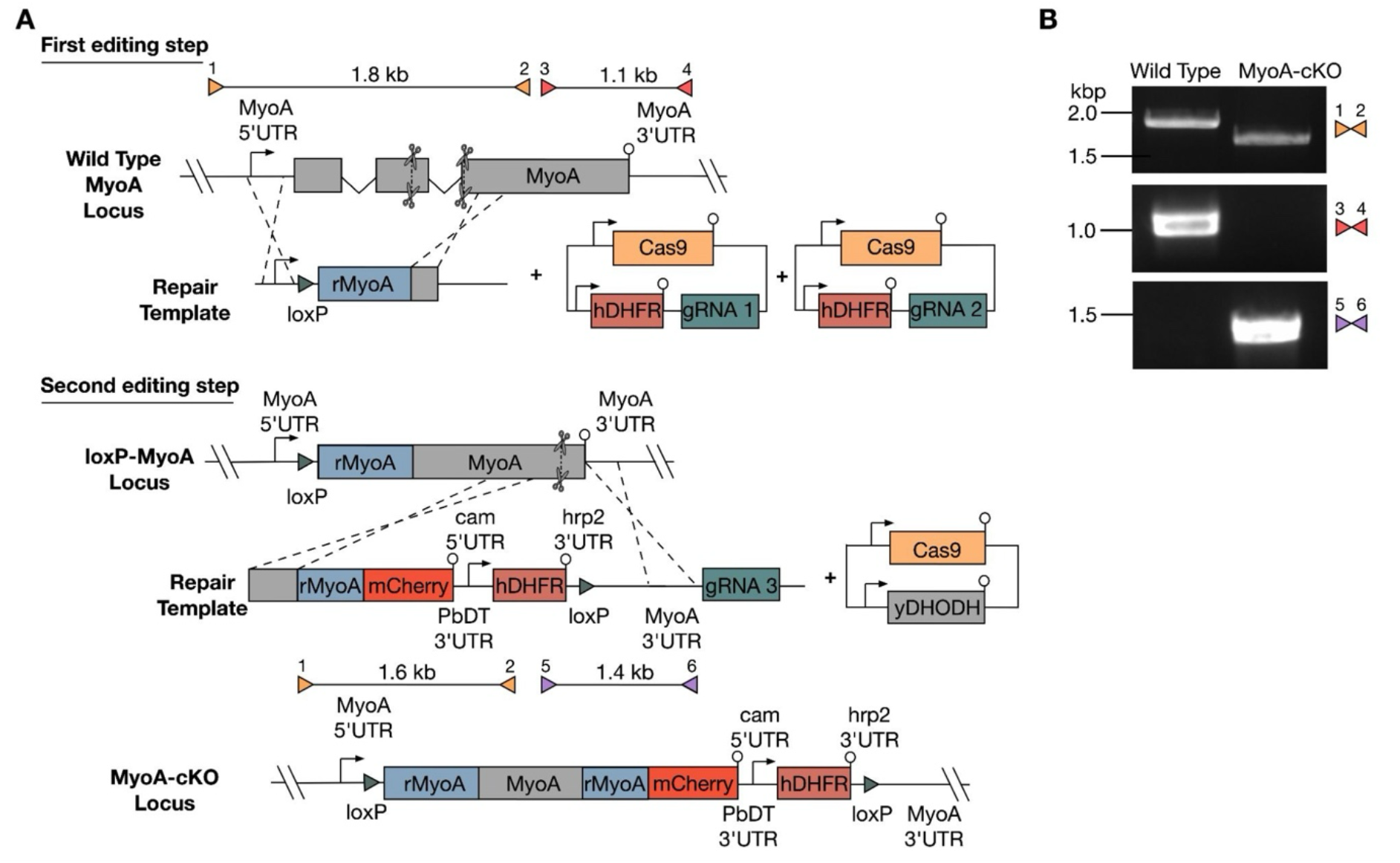
Generation of the MyoA-cKO line. **A** Schematic of the two-step CRISPR/Cas9-mediated approach used to generate the MyoA-cKO line. In the first editing step, a loxP site was introduced into the *myoa* promoter. In the second editing step, a C-terminal mCherry tag, a second loxP site and an hDHFR cassette were introduced at the endogenous *myoa* locus. Rapamycin treatment induces excision of the *myoA* gene between the two loxP sites. Scissors indicate CRISPR/Cas9 cleavage sites. Coloured triangles represent primer pairs used for diagnostic PCR screens together with the expected amplicon sizes. **B** Diagnostic PCR analysis of the parental WT and MyoA-cKO lines using the primer pairs shown in Supp.Fig2A, assessing loss of the WT locus (red triangles), integration of the promoter loxP site (orange triangles) and 5’ integration of the mCherry tag and second loxP site (purple triangles).

**Supplementary Table 1.**
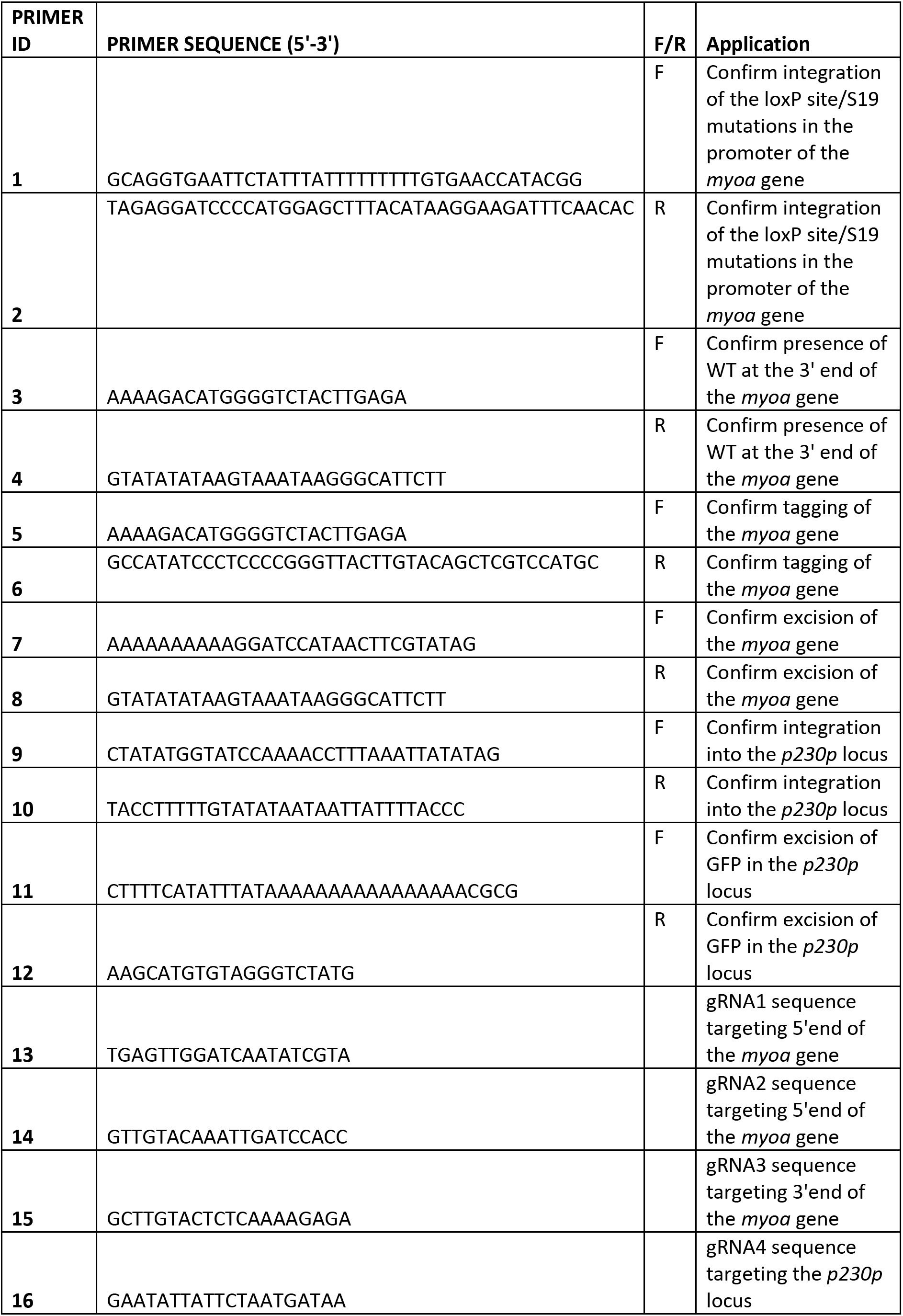

## Notes

### Competing Interest Statement

The authors have declared no competing interest.

